# Impact of 10-valent pneumococcal conjugate vaccine on invasive pneumococcal disease and nasopharyngeal carriage in Kenya

**DOI:** 10.1101/369876

**Authors:** Laura L. Hammitt, Anthony O. Etyang, Susan C. Morpeth, John Ojal, Alex Mutuku, Neema Mturi, Jennifer C. Moisi, Ifedayo M. Adetifa, Angela Karani, Donald O. Akech, Mark Otiende, Tahreni Bwanaali, Jackline Wafula, Christine Mataza, Edward Mumbo, Collins Tabu, Maria Deloria Knoll, Evasius Bauni, Kevin Marsh, Thomas N. Williams, Tatu Kamau, Shahnaaz K. Sharif, Orin S. Levine, J. Anthony G. Scott

**Author notes:** <u>Corresponding author</u>: Laura Hammitt, 415 N. Washington St., 4^th^ Floor Baltimore, MD 21231. <u>Alternate corresponding author:</u> Anthony Scott, PO Box 230, Kilifi, Kenya 80108.

## Abstract

**Background:** 10-valent pneumococcal conjugate vaccine (PCV10), delivered at 6, 10 and 14 weeks of age, was introduced in Kenya in January 2011, accompanied by a catch-up campaign in Kilifi County for children <5 years. Coverage with ≥2 PCV10 doses in children 2-11 months was 80% in 2011 and 84% in 2016; coverage with ≥1 dose in children 12-59 months was 66% and 87%, respectively.

**Methods:** Clinical and microbiological surveillance for invasive pneumococcal disease (IPD) among admissions of all ages at Kilifi County Hospital was linked to the Kilifi Health and Demographic Surveillance System from 1999-2016. We calculated the incidence rate ratio (IRR) comparing the pre-vaccine and post-vaccine eras, adjusted for confounding, and reported percent reduction in IPD as 1-IRR. Annual cross-sectional surveys of nasopharyngeal carriage were conducted from 2009-2016.

**Findings:** Surveillance identified 667 IPD cases in 3,211,403 person-years of observation. IPD incidence in children <5 years fell sharply in 2011 following PCV10 introduction, and remained low (PCV10-type IPD: 60·8 vs 3·2/100,000 [92% reduction; 95%CI: 78, 97]; overall IPD: 81·6 vs 15·3/100,000 [68% reduction; 95%CI: 40, 83]; 1999-2010 vs 2012-2016). PCV10-type IPD also declined significantly in unvaccinated age groups (<2 months, 5-14 years, ≥15 years), with estimated reductions of 100%, 74%, and 81%, respectively. There was no significant change in the incidence of non-PCV10 type IPD. In children aged <5 years, PCV10-type carriage declined by 74% and non-PCV10-type carriage increased by 71%.

**Interpretation:** Introduction of PCV10 in Kenya resulted in a substantial reduction in PCV10-type IPD in children and adults without significant replacement disease. These findings suggest that routine infant PCV10 immunization programmes with catch-up campaigns will provide substantial direct and indirect protection in low-income settings in tropical Africa.

## Background

The number of pneumococcal deaths in children aged 1-59 months was estimated at 332,000 globally in 2015, a decline of more than 50% from 2000.^1^ In middle- and high-income countries, inclusion of pneumococcal conjugate vaccines (PCVs) in routine infant vaccination programmes has led to a substantial reduction in the incidence of invasive pneumococcal disease (IPD) caused by vaccine serotypes (VT). In addition, because vaccinated children are less likely to carry and transmit VT pneumococci, programmatic use of PCV in children has resulted in a decline in IPD in unvaccinated individuals (i.e., “herd protection”).^2^ Evidence of impact of PCVs in low income settings is sparse and impact models rely upon efficacy estimates from randomized controlled trials, not real-world implementation. Furthermore, there is no evidence of the impact of 10-valent PCV conjugated to non-typeable *Haemophilus influenzae* (PCV10) on invasive pneumococcal disease in Africa, where the greatest burden of deaths from pneumococcal disease occur.

Sixty-one percent (82·4 million) of the world’s infants have not received PCV; however, dozens of low-income countries have recently introduced PCV or will do so in the coming decade.^3^ In 2011, with support from Gavi, the Vaccine Alliance, Kenya became one of the first countries in Africa to introduce PCV and the first country to use PCV10. PCV10 was introduced in the Kenyan national childhood immunization schedule as a 3-dose series administered at 6, 10, and 14 weeks of age. There is good evidence of the efficacy of PCV9 in African settings, but efficacy of PCV10 (Synflorix) has not been demonstrated.^4,5^ Furthermore, because of the potential for significant herd protection or for serotype replacement disease, the net population benefit of the PCV programme in these settings can only be estimated through longitudinal IPD surveillance. This information is essential to support realistic cost-effectiveness analyses and sustain the commitment of Ministries of Health (MOH) to the PCV programme as countries transition from Gavi support for vaccines to self-financing.

Through a collaboration between the Kenya MOH, Gavi, and the KEMRI-Wellcome Trust Research Programme (KWTRP), we used an existing integrated demographic, clinical, and microbiological surveillance system to conduct a prospectively designed assessment of vaccine impact against nasopharyngeal carriage and IPD in children and adults before and after introduction of PCV10 in the routine infant immunization programme in Kenya.

## Methods

### Setting

This study was conducted at KWTRP among residents of the Kilifi Health and Demographic Surveillance System (KHDSS), a rural community on the Kenyan coast covering an area of 891 km^2^. A census of the KHDSS in 2000 defined the resident population, and all subsequent births, deaths and migration events were monitored by fieldworker visits to every participating household at approximately 4-monthly intervals.^6^ The population (179568 in 1999; 239392 in 2007; 284826 in 2016) is served by a single government hospital, Kilifi County Hospital (KCH). Among women attending antenatal care at KCH, the prevalence of HIV infection ranged between 2·1% and 4·6% during 2005-2016 (Supplemental Figure 1). The prevalence of HIV among children <5 years in Kenya was estimated in 2012 at 1·6%.^7^ *Haemophilus influenzae* type b conjugate vaccine was introduced in Kenya in 2001.^8^ In the 12 years prior to PCV10 introduction, the serotypes contained in PCV10 (1, 4, 5, 6B, 7F, 9V, 14, 18C, 19F, and 23F) comprised 75% of IPD in children aged <5 years in Kilifi.

### Vaccine introduction and monitoring

In January 2011, the government of Kenya introduced PCV10 into the national immunization schedule, administered simultaneously with Pentavalent vaccine (diphtheria/whole cell pertussis/tetanus/hepatitis B/*H. influenzae* type b vaccine) at 6, 10 and 14 weeks of age. A national catch-up campaign provided three doses of PCV10 to children aged <12 months. As part of the study design, the MOH conducted a catch-up campaign in Kilifi County providing up to two doses of PCV10 to children aged 12-59 months in two campaigns, beginning on January 31, 2011 and March 21, 2011, each lasting 1-2 weeks. All vaccines were captured by the Kilifi Vaccine Monitoring System, a registry in which data clerks at 26 clinics serving the KHDSS link vaccination at the point of delivery to the child’s identification in the KHDSS.^9^ Coverage with PCV10 increased sharply during the catch-up campaign and slowly thereafter (Figure 1, Supplemental Table 1). Coverage with ≥2 PCV10 doses in 2-11 month olds was 80% in 2011 and 84% in 2016; coverage with ≥1 dose in 12-59 month olds was 66% and 87%, respectively.

**Figure 1.**
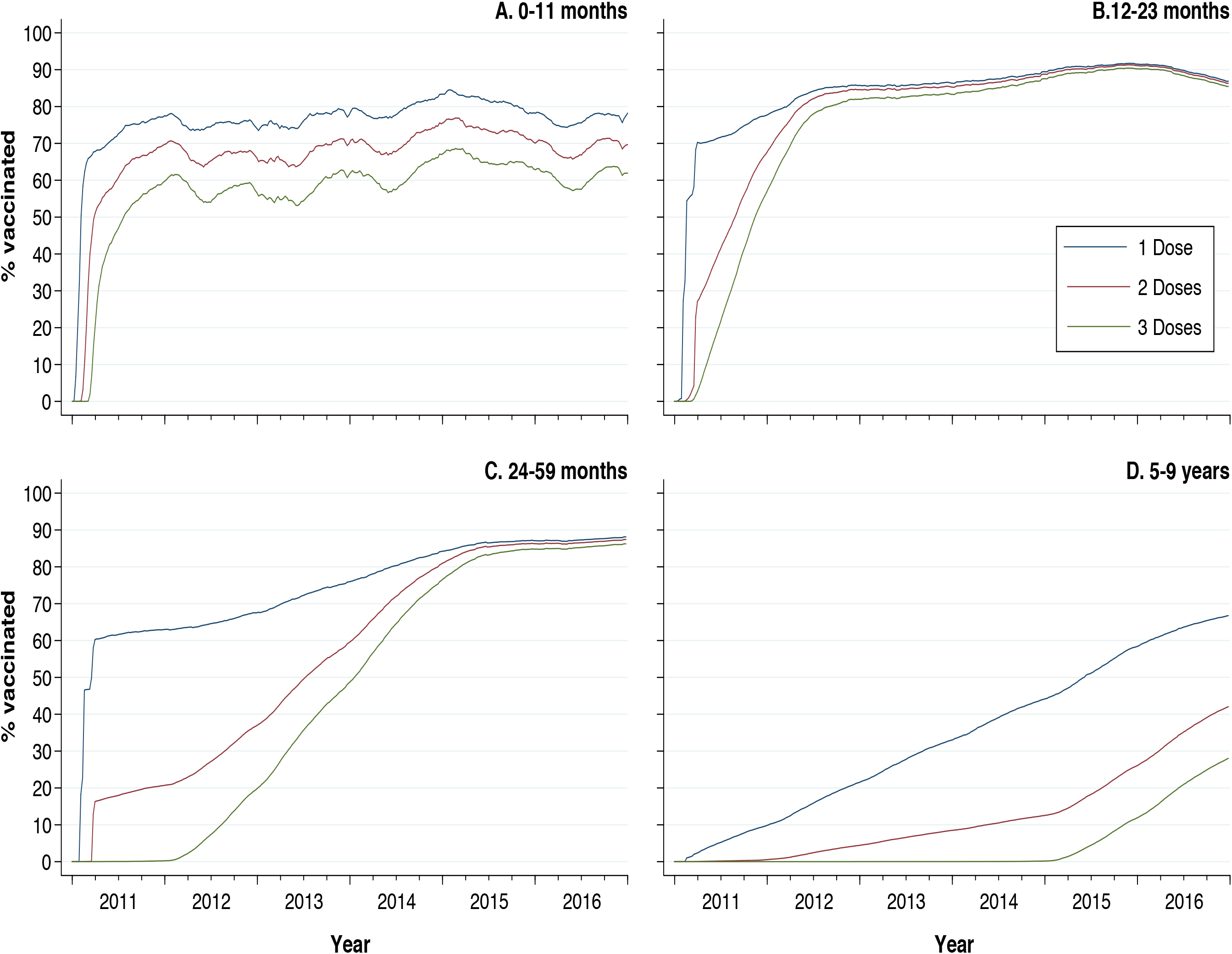
Proportion of children vaccinated with PCV10 (A: 0-11 months; B: 12-23 months; C: 24-59 months; C: 5-9 years) in the Kilifi Health and Demographic Surveillance System, 2011-2016.

### Clinical and laboratory procedures

Children admitted to KCH (with the exception of trauma patients or patients admitted for elective surgery) were investigated with a blood culture at the time of admission from 1999-2016.^10^ Adults (age ≥15 years) admitted to KCH from 2007-2016 were investigated with a blood culture at the time of admission if there were signs or symptoms of invasive bacterial disease (e.g., history of fever, axillary temperature <36·0°C or >37·5°C, signs of focal sepsis)^11^. Blood was cultured using an automated BACTEC instrument (BD Diagnostics, USA). From 1999-2016, with the exception of a brief change in practice in 2004-2005, the clinical indications for lumbar puncture were impaired consciousness or meningism in children younger than 5 years, prostration in children younger than 3 years, seizures (other than febrile seizures) in children younger than 2 years and suspicion of sepsis in children younger than 60 days, or suspected meningoencephalitis in adults. Cerebrospinal fluid (CSF) was cultured on horse blood and chocolate agar. Admitted patients were tested for HIV using two rapid antibody tests according to the Kenya national policy.^12^ Patients were treated according to Kenyan MOH or WHO guidelines.

Nasopharyngeal carriage of pneumococci was assessed through cross-sectional surveys conducted on annual samples of approximately 500 KHDSS residents of all ages selected at random from the KHDSS population register each year from 2009-2016. The methods are described elsewhere, with the exception that flocked swabs (Copan Diagnostics, USA) replaced rayon swabs in 2016.^13^

Isolates of *S. pneumoniae* from sterile-site and nasopharyngeal swab cultures were identified by optochin susceptibility; serotyping was performed by latex agglutination and Quellung reaction. If pneumococcal colonies of varying appearance were observed, only those of the dominant colony morphology were serotyped. Serogroup 6 isolates were tested by PCR for confirmation of serotype. Invasive isolates from 1999-2008 underwent repeat confirmatory serotyping by Quellung and multiplex PCR.^14^ Invasive isolates from 2008-2016 underwent real-time confirmatory serotyping by PCR; discordant results were resolved by a second PCR. A case of IPD was defined as isolation of *S. pneumoniae* from a sterile site culture in an individual admitted to KCH who was resident in the KHDSS. VT isolates were those belonging to PCV10 serotypes (1, 4, 5, 6B, 7F, 9V, 14, 19C, 19F, and 23F). All other serotypes were classified as non-VT. Pneumococcal meningitis was defined as isolation of *S. pneumoniae* from cerebrospinal fluid (CSF) or isolation of *S. pneumoniae* from blood, accompanied by a CSF white blood cell count of ≥50 x 10^6^ cells/L or a ratio of CSF glucose to plasma glucose of <0·1.^15^ Pneumococcal pneumonia was defined as a case of IPD in a child with cough or difficulty breathing, and ≥1 of the following: lower chest wall indrawing, central cyanosis, inability to drink, convulsions, lethargy, prostration, head nodding.^16^

### Statistical analysis

We designated January 1, 1999 through December 31, 2010, as the pre-vaccine era and January 1, 2012 through December 31, 2016, as the post-vaccine era. The year of vaccine introduction, 2011, was excluded from the analysis of impact. We calculated the age-stratified incidence of IPD in each year as the annual number of cases divided by the mid-year population in the KHDSS. We excluded admissions and person-years of observation during health care worker strikes (Supplemental Table 2). Unadjusted incidence rate ratios (IRRs) were calculated for the post-vaccine era compared to the pre-vaccine era by age group using negative binomial regression because of over-dispersion in the data. Possible confounders of the association between IPD and vaccine introduction included time (year), annual incidence of admissions, malaria admissions (i.e., presence of malaria parasites by microscopy), moderate or severe malnutrition admissions (among children <5 years; defined as weight-for-age >-2 z-scores below the median of the WHO child growth standards), and compliance with recommendations for investigation by blood culture. Potential confounders with a p-value <0·1 in univariate analysis were included in multivariable analysis; we used backward stepwise regression and excluded variables with a likelihood ratio test p-value ≥0·05. We built age-group specific models for overall IPD and applied the same structure within age-group for VT and non-VT IPD. The percent reduction in disease was calculated as 1 minus the adjusted IRR.

We calculated pneumococcal carriage prevalence ratios comparing nasopharyngeal carriage in the pre- and post-vaccine eras as previously described.^13^ Briefly, prevalence ratios were modeled using log-binomial regression; if the models failed to converge we used Poisson regression with robust confidence intervals. Adjusted prevalence ratios were age-standardized to reflect the stratified sampling scheme using the inverse of the sampling ratio as population weights.

The significance of vaccine impact on serotype-specific IPD or carriage was tested using a Bonferroni correction (i.e., for 25 serotypes, the correction was 0·05/25).

STATA 14·0 (Stata Corp, USA) was used for analysis.

### Ethical approval

The protocol was approved by the Oxford Tropical Ethical Review Committee (No. 30-10) and the Kenya National Ethical Review Committee (SSC1433). Adult participants and parents/guardians of all child participants provided written informed consent.

## Results

During the 18-year surveillance period, we identified 667 cases of IPD in 3,211,403 person years of observation among KHDSS residents (Supplemental Table 3). The specimen type and proportion of IPD cases with HIV infection did not change significantly between the pre- and post-vaccine eras (Table 1). Among children <5 years, the median age of IPD cases was 14 months (interquartile range [IQR] 7, 30) in the pre-vaccine era and 20 months (IQR 6, 38) in the PCV10 era. Throughout the 18-year surveillance period several indicators suggested marked improvements in the overall health of the population (Supplemental Figure 2; Supplemental Figure 3).

**Table 1.** Invasive pneumococcal disease among children and adults in the KHDSS admitted to the Kilifi County Hospital, in the pre-vaccine (1999-2010) and post-vaccine (2012-2016) eras.

|  | Pre-vaccine era<br>(1999-2010) |  | Post-vaccine era<br>(2012-2016) |  | p-value * |
| --- | --- | --- | --- | --- | --- |
|  | n | % | n | % |  |
| **Age <5 years** |  |  |  |  |  |
| Total admissions | 31,422 |  | 6,323 |  |  |
| Mean annual admissions | 2,618 |  | 1,264 |  | <0.001 |
| Recommended for blood culture | 30,098 | 96 | 5,691 | 90 | <0.001 |
| Blood culture collected† | 29,234 | 97 | 5,353 | 94 | <0.001 |
| <u>Culture-confirmed IPD</u> | 401 |  | 34 |  |  |
| Age <24 months | 273 | 68 | 20 | 59 | 0.27 |
| Male | 229 | 57 | 23 | 68 | 0.23 |
| Cultured from blood specimen | 390 | 97 | 34 | 100 | 0.33 |
| Cultured from CSF | 71 | 18 | 3 | 9 | 0.19 |
| Pneumococcal meningitis‡ | 78 | 62 | 3 | 38 | 0.16 |
| HIV-infected | 25 | 20 | 4 | 15 | 0.55 |
| Died during the episode | 70 | 17 | 11 | 32 | 0.03 |
| **Age 5-14 Years** |  |  |  |  |  |
| Total admissions | 5,296 |  | 2,060 |  |  |
| Mean annual admissions | 441 |  | 412 |  | 0.61 |
| Recommended for blood culture | 4,483 | 85 | 1,592 | 77 | <0.001 |
| Blood culture collected† | 4,200 | 94 | 1,438 | 90 | <0.001 |
| <u>Culture-confirmed IPD</u> | 127 |  | 22 |  |  |
| Male | 66 | 52 | 10 | 45 | 0.57 |
| Cultured from blood specimen | 121 | 95 | 21 | 95 | 0.97 |
| Cultured from CSF | 31 | 24 | 2 | 9 | 0.11 |
| Pneumococcal meningitis‡ | 37 | 76 | 4 | 67 | 0.64 |
| HIV-infected | 7 | 23 | 5 | 36 | 0.39 |
| Died during the episode | 30 | 24 | 2 | 9 | 0.13 |
| **Age ≥15 years** |  |  |  |  |  |
| Total admissions | 5,332 |  | 5,163 |  |  |
| Mean annual admissions | 1,326 |  | 1,032 |  | 0.10 |
| Recommended for blood culture | 5,048 | 95 | 4,597 | 89 | <0.001 |
| Blood culture collected† | 2,107 | 42 | 1,875 | 41 | 0.34 |
| <u>Culture-confirmed IPD</u> | 30 |  | 26 |  |  |
| Male | 12 | 40 | 11 | 42 | 0.86 |
| Cultured from blood specimen | 29 | 97 | 23 | 88 | 0.23 |
| HIV-infected | 4 | 33 | 14 | 66 | 0.09 |
| Died during the episode | 12 | 40 | 12 | 46 | 0.64 |

Abbreviations: IPD, invasive pneumococcal disease; CSF, cerebrospinal fluid
\*p-value computed by t-test for comparison of means and chi-square for comparison of percentages.
†Among those recommended for blood culture
‡Defined as isolation of *S. pneumoniae* from CSF or isolation of *S. pneumoniae* from blood, accompanied by a CSF white blood cell count of $50 \times 10^6$ cells/L or greater or a ratio of CSF glucose to plasma glucose less than 0.1.<sup>15</sup>

Among children <5 years, the incidence of VT-IPD declined from 60·8 to 3·2/100000 in the pre- and post-vaccine eras (Table 2; Figure 2; Supplemental Table 4) representing a reduction of 92% (95% CI: 78, 97; adjusted for year). The average annual number of VT-IPD cases fell from 25 (IQR 16, 33) in the pre-vaccine era to 1 (IQR 1, 2) in the post-vaccine era. Seven children had VT-IPD in the post-vaccine era: two were unvaccinated and five were age-appropriately vaccinated (Supplemental Table 5. Of the five children who developed VT-IPD after receipt of PCV10, two were noted to have underlying conditions (malnutrition in both). A decline in incidence was observed for all PCV10 serotypes, and this was statistically significant for serotypes 1 and 14, the most common serotypes in the pre-vaccine era. The incidence of non-VT IPD did not significantly increase following introduction of PCV10 (IRR 1·31; 95% CI: 0·65, 2·64). Overall, use of PCV10 led to a 68% (95% CI: 40, 83) reduction in the incidence of all-serotype IPD, and an 85% (95% CI: 66, 93) reduction in the incidence of bacteraemic pneumococcal pneumonia among children <5 years.

**Table 2.** Incidence (per 100,000) of invasive pneumococcal disease among children and adults in the KHDSS, in the pre- and post-vaccine eras.

| Serotype group and age category | Pre-vaccine era (1999-2010)* |  |  | Post-vaccine era (2012-2016) |  |  | Post- vs pre-vaccine era |  |  |  |
| --- | --- | --- | --- | --- | --- | --- | --- | --- | --- | --- |
|  | n | Incidence per 100,000 | 95% CI | n | Incidence per 100,000 | 95% CI | IRR† | 95% CI | Adjusted IRR‡ | 95% CI |
| **All-type IPD** |  |  |  |  |  |  |  |  |  |  |
| <2 months | 43 | 240.2 | 173.9, 323.6 | 1 | 7.9 | 0.2, 44.0 | 0.03 | 0.00, 0.25 | 0.13 | 0.01, 1.24 |
| <5 years | 401 | 81.6 | 73.8, 89.9 | 34 | 15.3 | 10.6, 21.4 | 0.18 | 0.11, 0.29 | 0.32 | 0.17, 0.60 |
| 5-14 years | 127 | 15.8 | 13.2, 18.8 | 22 | 5.5 | 3.5, 8.4 | 0.34 | 0.17, 0.67 | 0.47 | 0.24, 0.92 |
| ≥15 years | 30 | 2.4 | 1.6, 3.4 | 26 | 3.4 | 2.5, 5.7 | 0.56 | 0.33, 0.96 | 0.63 | 0.36, 1.08 |
| **PCV10-type** |  |  |  |  |  |  |  |  |  |  |
| <2 months | 31 | 173.2 | 117.7, 245.8 | 0 | 0.0 |  | Not estimable |  |  |  |
| <5 years | 299 | 60.8 | 54.1, 68.1 | 7 | 3.2 | 1.3, 6.5 | 0.05 | 0.02, 0.12 | 0.08 | 0.03, 0.22 |
| 5-14 years | 105 | 13.1 | 10.7, 15.8 | 10 | 2.5 | 1.2, 4.6 | 0.19 | 0.08, 0.43 | 0.26 | 0.11, 0.59 |
| ≥15 years | 20 | 1.6 | 1.0, 2.4 | 5 | 0.8 | 0.2, 1.7 | 0.16 | 0.06, 0.45 | 0.19 | 0.07, 0.51 |
| **Non-PCV10 type** |  |  |  |  |  |  |  |  |  |  |
| <2 months | 12 | 67.0 | 34.3, 117.0 | 1 | 7.9 | 0.2, 4.0 | 0.12 | 0.02, 0.94 | 0.26 | 0.02, 3.45 |
| <5 years | 102 | 20.8 | 16.9, 25.2 | 27 | 12.2 | 8.0, 17.7 | 0.58 | 0.35, 0.95 | 1.31 | 0.65, 2.64 |
| 5-14 years | 22 | 2.7 | 1.7, 4.2 | 12 | 3.0 | 1.6, 5.3 | 1.10 | 0.54, 2.22 | 1.45 | 0.66, 3.20 |
| ≥15 years | 10 | 0.8 | 0.4, 1.5 | 21 | 3.1 | 1.9, 4.8 | 1.36 | 0.62, 2.98 | 1.47 | 0.67, 3.21 |
| **All-type pneumococcal pneumonia** |  |  |  |  |  |  |  |  |  |  |
| <5 years | 212 | 43.1 | 37.5, 49.3 | 19 | 8.6 | 5.2, 13.4 | 0.20 | 0.11, 0.35 | 0.15 | 0.07, 0.34 |
| 5-14 years | 57 | 7.1 | 5.4, 9.2 | 11 | 2.8 | 1.4, 5.0 | 0.38 | 0.17, 0.86 | 0.49 | 0.21, 1.11 |
| **All-type pneumococcal meningitis** |  |  |  |  |  |  |  |  |  |  |
| <5 years | 78 | 15.9 | 12.5, 19.8 | 3 | 1.4 | 0.3, 4.0 | 0.08 | 0.02, 0.29 | 0.31 | 0.08, 1.21 |
| 5-14 years | 37 | 4.6 | 3.2, 6.4 | 4 | 1.0 | 0.3, 2.6 | 0.21 | 0.06, 0.71 | 0.31 | 0.09, 1.00 |

Abbreviations: CI, confidence interval; IRR, incidence rate ratio
\*For individuals ≥15 years, the pre-vaccine era was 2007-2010.
†IRR estimated using negative binomial regression.
‡IRR estimated using negative binomial regression, adjusted for confounding factors significant in the age-specific all-type IPD models: year (age groups <2 months and <5 years), blood culture collection (age groups 5-14 years and ≥15 years).

**Figure 2.**
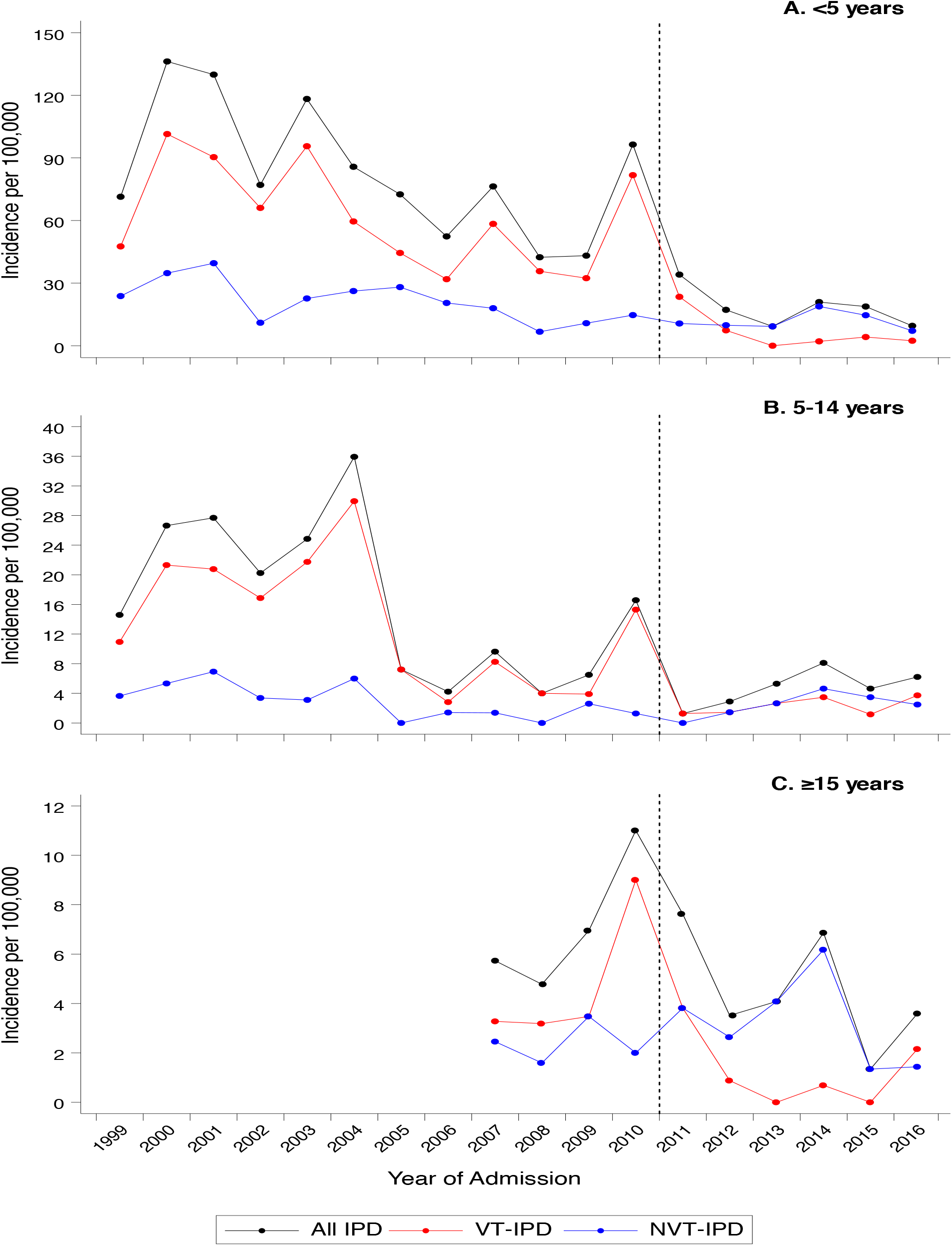
Incidence of overall (black line), vaccine-type (red line) and non-vaccine type (blue line) invasive pneumococcal disease (IPD) in individuals aged A) <5 years; B) 5-14 years; C) ≥15 years in the Kilifi Health and Demographic Surveillance System, 1999-2016. Vertical dashed line indicates PCV10 introduction.

Among children too young to be vaccinated (i.e., <2 months of age), the incidence of VT-IPD declined from 173·2 to 0/100000. A significant decline in the rate of VT-IPD was also observed among individuals 5-14 years (74% reduction, 95% CI: 41, 89; adjusted for blood culture ascertainment), and ≥15 years (81% reduction, 95% CI: 49, 93; adjusted for blood culture ascertainment) (Figure 2; Table 2, Supplemental Table 3). There was no significant change in the incidence of non-VT IPD in these age groups (Table 2).

Overall 4,066 KHDSS residents were enrolled in the nasopharyngeal carriage surveys (Supplemental Table 6). VT carriage declined among individuals <5 years (with statistically significant reductions for serotypes 6B, 19F and 23F), 5-14 years, and ≥15 years. Non-VT carriage increased significantly in in all age groups (Table 3; Supplemental Figure 4). Among children <5 years, carriage of vaccine-related serotype 19A increased; no effect on carriage of serotype 6A was observed. In 2016, carriage of VT pneumococci was detected in 10/156 (6%) of children <5 years, all of whom had received 3 doses of PCV10, and 7/99 (7%) of children 5-14 years, none of whom had received PCV10.

**Table 3.** Carriage prevalence and prevalence ratios for nasopharyngeal carriage of *Streptococcus pneumoniae* and non-typeable *Haemophilus influenzae* in the pre- and post-vaccine eras.

|  | Carriage prevalence<br>Pre-vaccine era<br>(2009-2010) |  | Carriage prevalence<br>Post-vaccine era<br>(2012-2016) |  | Crude prevalence<br>ratio (PR) |  | Age-standardized<br>adjusted prevalence<br>ratio (aPR)* |  |
| --- | --- | --- | --- | --- | --- | --- | --- | --- |
|  | n | % | n | % | PR | 95% CI | aPR | 95% CI |
| *All S. pneumoniae* |  |  |  |  |  |  |  |  |
| <5 years | 229 | 74.4 | 606 | 76.0 | 1.02 | 0.95, 1.10 | 1.00 | 0.92, 1.08 |
| 5-14 years | 103 | 52.6 | 237 | 48.5 | 0.92 | 0.78, 1.08 | 0.96 | 0.81, 1.13 |
| ≥15 years | 123 | 24.0 | 287 | 22.8 | 0.95 | 0.79, 1.14 | 1.00 | 0.81, 1.22 |
| *Vaccine-type S. pneumoniae* |  |  |  |  |  |  |  |  |
| <5 years | 104 | 33.8 | 70 | 8.8 | 0.26 | 0.20, 0.34 | 0.26 | 0.19, 0.35 |
| 5-14 years | 30 | 15.3 | 29 | 5.9 | 0.39 | 0.24, 0.63 | 0.38 | 0.22, 0.64 |
| ≥15 years | 29 | 5.7 | 17 | 1.4 | 0.24 | 0.13, 0.43 | 0.23 | 0.12, 0.44 |
| *Non vaccine-type S. pneumoniae* |  |  |  |  |  |  |  |  |
| <5 years | 125 | 40.6 | 557 | 69.9 | 1.72 | 1.49, 1.99 | 1.71 | 1.47, 1.99 |
| 5-14 years | 73 | 37.2 | 216 | 44.2 | 1.19 | 0.96, 1.46 | 1.25 | 1.01, 1.54 |
| ≥15 years | 94 | 18.3 | 286 | 22.7 | 1.24 | 1.01, 1.53 | 1.30 | 1.05, 1.63 |

Abbreviations: CI, confidence interval
\* Models adjusted for number of children aged <10 years in household, month of swab collection, cough or rhinorrhea in preceding 14 days (all age groups); antibiotic use in the preceding 14 days also remained in the model for children age 5-14 years.

## Discussion

Using a longstanding, integrated clinical, laboratory and demographic surveillance system, we documented a 92% reduction in VT-IPD in children aged <5 years and substantial indirect protection in Kilifi, Kenya following introduction of PCV10 to the routine infant immunization schedule, accompanied by a catch-up campaign. Kenya was the first African country to include PCV10 in its routine childhood immunization programme. This study provides the first population-level evidence of a robust direct and indirect effect of a PCV10 programme in a low-income country and does not find significant evidence of serotype replacement disease in the first six years of PCV10 use.

Following introduction of PCVs, a large decline in IPD was documented in numerous developed world settings; however, serotype distributions in carriage and IPD differ by geographic area and there have been few opportunities to examine PCV10 impact in a developing country.^17^ In Kilifi, we observed a 68% reduction in overall IPD and a 92% reduction in VT-IPD among children <5 years, consistent with findings in middle- and high-income settings using either PCV10 or PCV13. Four years after PCV13 introduction, IPD was reduced by 64% in US children <5 years and 46% in British children <2 years.^18,19^ Within the first 3-5 years of PCV13 use, the rate of IPD caused by the 6 serotypes present in PCV13 but not PCV7 declined 93% in US children <5 years, 80% in Alaska Native children <5 years, 89% in British children <2 years, and 82% in Gambian children 2-23 months.^18-21^ In Latin American countries using PCV10 or PCV13, effectiveness against VT-IPD has been estimated at 56% to 84%.^22^ Contributing to the observed reduction in IPD in Kilifi were an 85% reduction in bacteraemic pneumococcal pneumonia incidence and a 69% reduction in pneumococcal meningitis incidence in children <5 years. PCV impact on these important clinical outcomes has been documented elsewhere.^23-25^ Given that the majority of pneumococcal disease is comprised of pneumonia, it is notable that PCV10 introduction in Kenya was associated with reductions in childhood hospitalisations with clinically-defined and radiologically-confirmed pneumonia of 27% and 48%, respectively.^26^

In the pre-vaccine era, IPD was driven by epidemics of serotypes 1 and 5 in Kilifi. PCV10 use not only reduced the incidence of disease but obliterated IPD epidemics, as was also observed in the US.^27^ We did not observe a reduction in IPD caused by vaccine-related serotypes 6A or 19A; this is consistent with the findings from the nasopharyngeal carriage surveys in Kilifi. In contrast, a recent analysis of the long-term impact of PCV10 in Finland documented a reduction in IPD caused by serotype 6A (but not 19A).^28^

Notably, we observed protection among infants too young to be vaccinated and among older children and adults. This contrasts with findings from The Gambia, where indirect effects were not observed.^21^ A catch-up campaign was not undertaken in The Gambia but in Kilifi this likely accelerated population protection.^29^ In South Africa a decline in VT-IPD was noted among adults aged 25-44 years within 3 years of PCV7 introduction.^30^

The indirect protection afforded by PCVs is driven by the reduction in nasopharyngeal carriage of VT pneumococci among vaccinated children. In Kilifi, there was a significant reduction in carriage of VT pneumococci in both vaccinated and unvaccinated populations within 6 months of PCV10 introduction.^13^ However, while vaccine-type carriage has declined, VT pneumococci continue to be identified in 6% of children <5 years and 8% of infants in Kilifi, compared to <1-2% in other countries that use PCVs.^31-35^ This may reflect a higher force of infection in Kilifi and it indicates continued risk for VT-IPD in unvaccinated or under-vaccinated children, and adults.^36^ An alternative explanation for the persistence of VT carriage in Kilifi is that, unlike most middle-income and high-income countries, Kenya introduced PCV10 without a booster dose in the second year of life. Many low-income countries have introduced PCV with three primary doses without a booster (3+0 schedule), and it will be important to determine whether the absence of a booster dose, which may be accompanied by waning immunity, leads to a persistent transmission reservoir, vaccination failures or rebound disease incidence.

In addition to persistent VT carriage, we also observed a 71% increase in carriage of non-VT pneumococci (particularly serotype 19A) in children <5 years. The PCV-associated decline in VT carriage and corresponding increase in non-VT carriage has been well-described. Although an increase in non-VT IPD disease has been reported in among some age groups in several settings that use expanded valency PCVs, the increases have been small compared to the decline in VT-IPD.^18,19^ The only exception to this is a recent increase in total IPD incidence among persons 5-64 and ≥65 years of age in North East England that was observed seven years following introduction of PCV13 in the routine vaccination program in the United Kingdom.^37^ Although our surveillance for non-VT IPD did not detect a significant increase in any age group, the direction of change was positive in all age-groups ≥2 months (IRRs 1·31-1·47). Given the low baseline incidence of non-VT IPD, comprising only one quarter of the pre-vaccine burden of IPD among children <5 years, the power of the study was only sufficient (i.e., >80%) to detect a ≥2·3-fold change (Supplemental Table 7). The small relative increase that was observed did not translate into a significant absolute rise in incidence. To clarify these emerging trends, it will be important to continue to monitor children and adults for pneumococcal disease in the present surveillance setting for several years to come.

The before-after study design, which is the principal method for evaluation of population impact of vaccines, has inherent weaknesses. In Kilifi, general health improved slowly over the surveillance period. The incidence of hospital admissions for a variety of illnesses declined. The reduction in non-VT IPD over time among infants <2 months, in contrast to the rise in older age groups, suggests specific improvements in maternity services and infant care. In an exploratory post-hoc analysis using negative binomial regression, adjusting for calendar-year, the introduction of PCV10 was associated with a non-statistically significant 35% reduction in invasive *S. aureus* disease in children <5 years (IRR 0·65; 95% CI: 0·36, 1·18). The incidence of HIV in the population has not been systematically measured and was not included in the analysis; however, the prevalence of HIV among women seeking ante-natal care was <5% over the surveillance period and it is therefore unlikely that changes in HIV incidence would have significantly confounded estimates of vaccine effectiveness. Overall, several factors argue that the observed reduction in IPD in Kilifi is attributable to the introduction of PCV10. Consistent surveillance methods were used over a long period of time and changes in VT-IPD occurred abruptly at the same time as vaccine introduction with a catch-up campaign, and simultaneously with marked changes in VT carriage prevalence. Systematically collected data on a wide range of possible confounders were included in the analysis.

Pneumococcal disease remains a leading vaccine-preventable cause of childhood mortality, and the majority of these deaths occur in Africa. To date, 141 countries, including 58 Gavi-eligible countries, have introduced a PCV into their national childhood immunization programmes.^3^ Currently, sixteen countries were in the process of transitioning out of Gavi support while five had reached the end of Gavi support. The PCV programme is the most expensive component of the national immunization schedule and the sustainability of PCV vaccination in low-income countries will depend on the demonstrable impact of PCV in reducing childhood morbidity and mortality. Because of the necessity for stable pre-vaccine surveillance, evaluations of PCV impact are rare in Africa. Based on this carefully standardized, prospectively designed, 18-year surveillance study, we conclude that use of PCV10 in tropical Africa will lead to substantial and sustained health benefits for the whole population.

## Acknowledgements

We thank the residents of the KHDSS, the Ministry of Health Sub-County Health Management Team in Kilifi County, and the dedicated team of field workers, administrative staff, clinicians and laboratorians who worked on this study. This paper is published with the permission of the Director, Kenya Medical Research Institute.

**Funding:** Gavi, The Vaccine Alliance; The Wellcome Trust of Great Britain

